# AgroGem: A Rapid and Scalable Transient Transformation System for Functional Genetics in Multiple Plant Species

**DOI:** 10.64898/2026.07.03.736435

**Authors:** Shengsong Guo, Oliver Schlegel, Jitesh Kumar, Zachary Myers, Shahryar Kianian, Kathleen Greenham, Feng Zhang

## Abstract

Plant genetic transformation technologies are essential for functional genomics and genome engineering in plants. While transient expression systems offer a rapid alternative to stable transformation, existing platforms are often constrained by low efficiency, technical complexity, and limited scalability. Here, we developed AgroGem, an efficient Agrobacterium-mediated transient transformation system utilizing a geminiviral replicon-based T-DNA vector for Arabidopsis and *Brassicaceae* species. AgroGem significantly outperformed existing transient approaches, including AGROBEST and protoplast-based assays, in CRISPR-mediated editing efficiency. Moreover, AgroGem recapitulated the mutation spectra and chromatin accessibility-dependent editing patterns observed in stable transformation across both Cas9 and Cas12a systems, indicating that it captures genome editing outcomes in native chromatin contexts. Leveraging this capability, we performed high-resolution profiling of CRISPR-induced mutation outcomes across a panel of DNA repair mutants and identified distinct repair signatures, including unexpected roles for KU80 and XRCC4 in regulating non-homologous end joining (NHEJ). AgroGem also supported bimolecular fluorescence complementation assays for protein-protein interaction studies in Arabidopsis and was readily adapted to plate-based formats for high-throughput applications. Together, these results establish AgroGem as a robust, scalable, and versatile platform for genome editing, DNA repair analysis, and functional genetics in plants.

## Introduction

Plant genetic transformation technologies are essential for characterizing gene function and for advancing genome engineering (Altpeter et al., 2016). In plants, transformation is generally achieved through either stable or transient approaches. Stable transformation results in genomic integration of transgenes with heritable gene expression across generations, but it is time-consuming and often constrained by species and genotype dependence (Goralogia et al., 2025). In contrast, transient transformation provides the means for short-term transgene expression without requiring stable integration (Yoo et al., 2007; Cardi et al., 2023). While stable transformation is needed for generating transgenic lines and assessing long-term phenotypes, transient transformation systems are routinely used for rapid evaluation of gene function, and more recently for testing genome editing reagents in plants (Lin et al., 2018; Chen et al., 2019).

Several transient transformation systems have been developed in plants and can be broadly grouped into two categories: *Agrobacterium*-mediated and direct delivery approaches. *Agrobacterium*-mediated methods introduce transgenes via T-DNA transfer and are generally simple and inexpensive (Wroblewski et al., 2005; Wu et al., 2014; Nasti et al., 2021). However, their performance is often highly species dependent and can be limited by variable transformation efficiency. For example, leaf agroinfiltration is routinely robust in *Nicotiana benthamiana* but remains less robust in other plant species, including *Arabidopsis thaliana*, even though *Agrobacterium*-mediated stable transformation has been well established in this model system (Wroblewski et al., 2005; Wu et al., 2014).

In contrast, for species, tissues, or genotypes that are difficult to transform using *Agrobacterium*, direct delivery approaches such as protoplast transfection and particle bombardment have been used (Yoo et al., 2007; Gunadi et al., 2019). However, protoplast-based assays require enzymatic cell wall digestion and specialized handling, often yield variable transformation efficiency, and introduce stress responses that complicate downstream analysis. Moreover, both approaches can be challenging to scale up for high-throughput formats (Yoo et al., 2007; Lin et al., 2018). These limitations highlight the need for a simple, robust, and scalable transient transformation system that can be readily implemented across plant species.

Development of scalable transient transformation systems in model plants such as Arabidopsis is particularly attractive given the extensive genetic and genomic resources available. In Arabidopsis, the protoplast transfection approach was first developed for rapid gene function and genome editing studies (Yoo et al., 2007; Banakar et al., 2022). However, even though it can be adapted to plate-based, high-throughput formats, preparation of high-quality protoplasts is technically demanding, expensive, and time-intensive, and the stress response remains another concern (Banakar et al., 2022). In parallel, several *Agrobacterium*-mediated transient expression methods, including AGROBEST, Fast-TrACC, and Agroflood, have been developed for gene function studies (Wu et al., 2014; Nasti et al., 2021; Nagalakshmi et al., 2022). Despite their utility, *Agrobacterium*-mediated approaches frequently exhibit low and inconsistent transformation efficiency, making them less suitable for quantitative comparisons or systematic screening. There is therefore a clear need for a transient transformation platform that is robust, scalable, and compatible with high-throughput functional genetics. An ideal system should combine high transformation efficiency with a simple workflow, rapid turnaround time, and compatibility with plate-based formats. Importantly, such a system should preserve endogenous biological contexts and physiological conditions to enable robust evaluation of gene function and genome editing outcomes.

In this study, we report the development of AgroGem, an improved *Agrobacterium*-mediated transient expression system optimized for Arabidopsis and transferable to related species. By optimizing *Agrobacterium* strain selection, T-DNA backbone design, and co-cultivation conditions, we establish a scalable and efficient transient transformation method. We demonstrate that AgroGem enables rapid and quantitative evaluation of CRISPR-based genome editing systems, preserves chromatin context-dependent editing effects, and supports high-resolution genetic dissection of DNA repair pathways. In addition, we show that AgroGem can be applied to protein-protein interaction assays and extended to multiple *Brassicaceae* species in a plate-based format. Together, these results establish AgroGem as a versatile and scalable platform for functional genetics and genome editing in plants.

## Results

### Optimization of Agrobacterium-mediated transient expression system in Arabidopsis

To develop a scalable *Agrobacterium*-mediated transient transformation method in *Arabidopsis thaliana*, we chose to adopt the AGROBEST method because of its simplicity and potential compatibility with plate-based assays (Wu et al., 2014). One- or Two-week-old Arabidopsis seedlings (Col-0) were co-cultivated with the *Agrobacterium tumefaciens*, which carried a T-DNA construct with the fluorescent reporter, AmCyan, driven by *Solanum lycopersicum* ubiquitin (*Sl*Ubi) promoter. We first tested three commercially available *Agrobacterium tumefaciens* strains, C58C1, GV3101, and the hypervirulent strain EHA105, under a 22°C 16-hour light/8-hour dark photoperiod for three days, following the previously described AGROBEST conditions with either one- or two-week old seedlings (Koncz and Schell, 1986; Hood et al., 1993; Gelvin, 2003). Under these conditions, no visible fluorescence signal was observed for any of the strains tested using the wildtype *Arabidopsis* Col-0 background (Figure 1A; only 2-week-old seedlings were shown).

**Figure 1.**
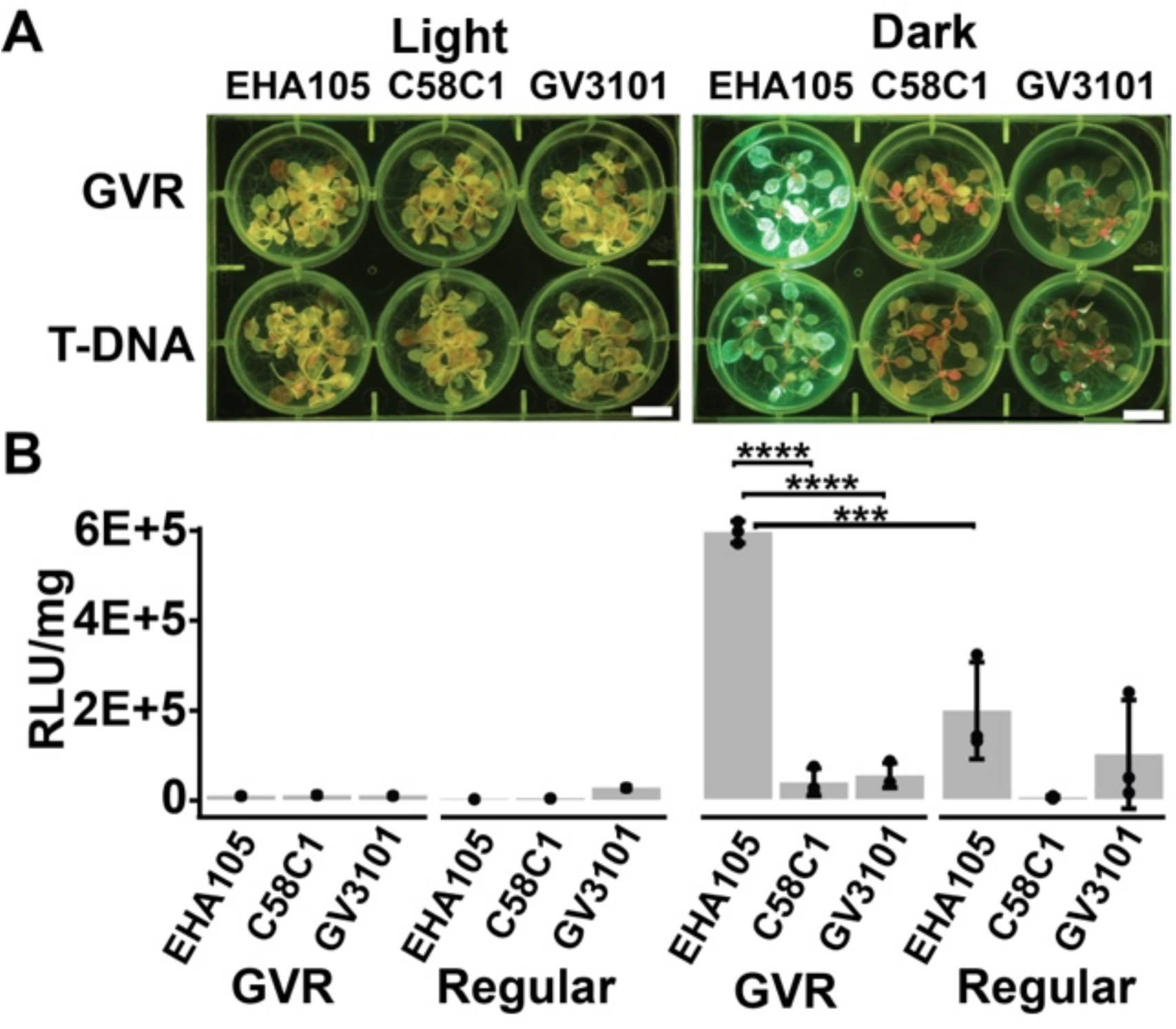
Optimization of Agrobacterium-mediated transient transformation in Arabidopsis. (A) Evaluation of Agrobacterium strain, T-DNA vector design, and co-cultivation conditions for transient expression in 2-week-old Arabidopsis seedlings. Three commercially available Agrobacterium tumefaciens strains with different virulence backgrounds, EHA105, C58C1, and GV3101, were tested in combination with either a geminiviral replicon T-DNA construct (GVR) or a conventional T-DNA construct lacking viral replication elements (Regular). Seedlings were co-cultivated under either standard long-day light conditions (22°C, 16 h light/8 h dark; “Light”) or continuous darkness for 3 days (22°C, “Dark”). Representative AmCyan fluorescence images are shown. Scale bars: 1 cm. (B) Quantification of transient gene expression by luciferase assay for each treatment is shown in (A). Luciferase activity was measured from independently transformed samples with the relative light unit (RLU) normalized to the fresh weight of each sample (mg). Three biological replicates were performed for each assay (*** *p* value < 0.001; **** *p* value < 0.0001; error bars are standard deviation). The source data are provided in the Supplemental Source Data file S1.

To determine whether transgene expression contributed to the low signal, we replaced the conventional T-DNA backbone with a Bean yellow dwarf virus (BeYDV) derived geminiviral construct, which undergoes rolling-circle replication and is expected to increase transgene copy number and expression levels (Baltes et al., 2014; Supplemental Figure S1). However, this change did not result in a noticeable improvement in fluorescence intensity under the same co-cultivation conditions.

Agrobacterium has been reported to exhibit increased motility, enhanced adhesion to plant tissues, and elevated virulence under dark conditions, leading to more efficient infection (Oberpichler et al., 2008). We therefore performed co-cultivation in complete darkness for 3 days. Under these conditions, the combination of the geminiviral backbone and the EHA105 strain produced the highest AmCyan fluorescence level (Figure 1A). The conventional T-DNA backbone also yielded detectable fluorescence, although at substantially lower intensity than the geminiviral construct (Figure 1A). In contrast, GV3101 produced a much weaker signal, while C58C1 exhibited little to no detectable fluorescence (Figure 1A).

To further quantify transformation efficiency, we incorporated a luciferase reporter driven by the Arabidopsis *Ubiquitin 10* (*AtUbi10*) promoter into the transformation construct. Luciferase activity showed a strong correlation with the observed AmCyan fluorescence intensity (Figure 1B). Across all conditions tested, the combination of the geminiviral backbone, the EHA105 strain, and dark co-cultivation resulted in the highest luciferase expression levels. Although the conventional T-DNA backbone was also capable of mediating transient transgene delivery under these conditions, it produced approximately one-third of the luciferase activity observed with the geminiviral construct. Overall, dark co-cultivation substantially enhanced transformation efficiency relative to light-grown conditions when using the EHA105 strain (Figure 1B). We termed this optimized method AgroGem, an improved Agrobacterium-mediated transient expression system that incorporates the geminiviral replicon vector and optimized co-cultivation conditions to achieve efficient transgene delivery and expression in Arabidopsis.

### AgroGem enables rapid evaluation of genome editing systems

Following the optimization of the AgroGem transient expression system, we next evaluated whether this platform could be used for rapid and quantitative assessment of CRISPR-based genome editing systems, which are most commonly analyzed using protoplast-based assays (Chen et al., 2019). We first compared CRISPR-Cas9 editing efficiencies across three transient expression approaches, AgroGem, AGROBEST, and protoplast-based methods, by targeting the endogenous *magnesium chelatase I2* (*CHLI2)* site, as described previously (Weiss et al., 2022). Genomic DNA was extracted from individual samples, followed by PCR amplification of the target site and next-generation sequencing to quantify editing frequencies and repair outcomes. AgroGem achieved the highest average editing frequency (21.14%), representing a 10.84-fold and 2.98-fold increase relative to AGROBEST (1.95%) and protoplast-based methods (7.08%), respectively (Figure 2A).

**Figure 2.**
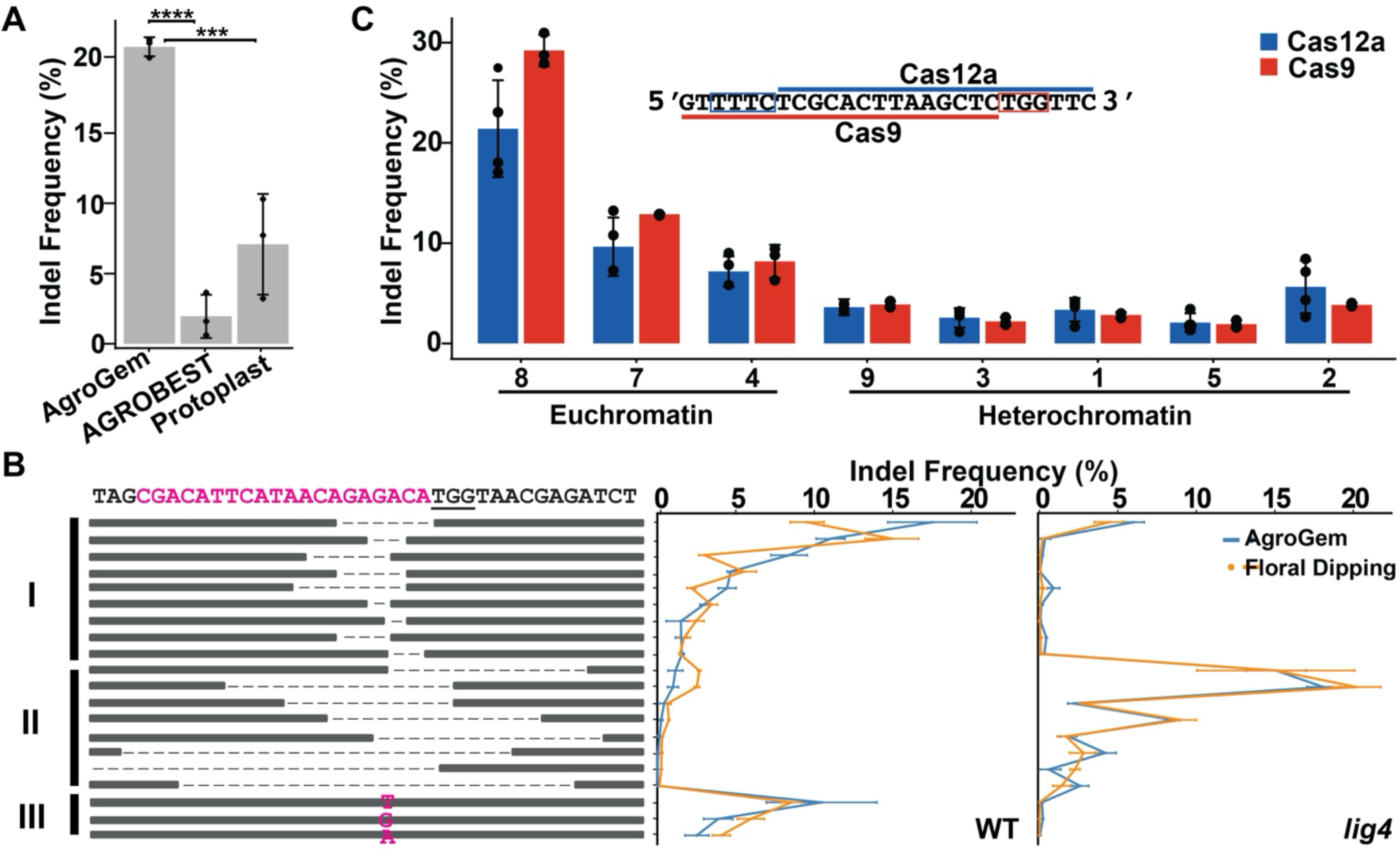
AgroGem supports efficient genome editing and preserves chromatin-dependent editing outcomes. (A) Comparison of CRISPR–Cas9 editing efficiencies at the *CHLI2* site using three transient expression systems: AgroGem, AGROBEST, and protoplast transfection. Indel frequencies were quantified by next-generation sequencing. Three biological replicates were performed for each test. (B) Relative indel frequency distributions generated by AgroGem (blue) and stable floral dip transformation (orange) at the *CHLI2* site in wild type and the *lig4* mutant backgrounds. Top 20 mutation types are shown with average relative frequencies from three biological replicates. In deletion classes (Class I: deletion size < 10 bp; Class II: deletion size > 10 bp), dashed lines indicate deleted nucleotides, while in insertion classes (Class III: 1 bp insertions), individual nucleotides denote the identity of the inserted base. (C) Chromatin accessibility influences editing efficiency for both *Lb*Cas12a (blue) and *Sp*Cas9 (red). Indel frequencies were measured at multi-copy target sites located in euchromatin or heterochromatin. The target sequence (indicated with underlines) and protospacer adjacent motifs (PAM; indicated by boxes) for *Lb*Cas12a (blue) and *Sp*Cas9 (red) are indicated above. Three biological replicates were performed for the *Sp*Cas9 assay, and four for the *Lb*Cas12a assay (*** *p* value < 0.001; **** *p* value < 0.0001; error bars are standard deviation). The source data are provided in the Supplemental Source Data file S1.

While editing frequency provides an important measure of performance, mutation spectra offer deeper insight into DNA repair outcomes. We therefore examined whether the mutation profiles generated using the AgroGem approach were comparable to those produced by stable transformation via the floral dip method, as reported previously (Weiss et al., 2022; Weiss et al., 2024). Insertion and deletion (indel) profiles at the *CHLI2* site were analyzed across three genetic backgrounds: wild type (Col-0) and the *lig4* knockout mutant defective in DNA end ligation during canonical non-homologous end joining (NHEJ) (Qi et al., 2013). In the wild-type background, CRISPR-induced mutation profiles were highly similar between AgroGem and floral dip transformation, with a Pearson correlation coefficient of 0.83 (*p* value = 5.73 x 10^-6^), and were dominated by 1-10 bp indels (Figure 2B). Similarly, mutation profiles in the *lig4* mutant backgrounds were highly correlated between the two approaches, with Pearson correlation coefficients of 0.99 (*p* value = 1.66 x 10^-16^; Figure 2B). Notably, indel distributions in the *lig4* mutant backgrounds were shifted toward larger deletions, consistent with previous observations (Figure 2B; Qi et al., 2013). Together, these results demonstrate that AgroGem recapitulates the mutation spectra generated by stable transformation, supporting its use as a rapid platform for evaluating genome editing outcomes.

Previous studies have shown that CRISPR-Cas9 editing efficiencies are strongly influenced by chromatin context when assessed using stable floral dip transformation. To determine whether the AgroGem approach preserves these chromatin-dependent effects, we analyzed editing rates at a previously characterized multi-copy CRISPR site (MC site) repeated at eight loci spanning diverse chromatin environments (Weiss et al., 2022). Consistent with results obtained using stable transgenic approaches, Cas9-induced mutagenesis rates measured with AgroGem followed the same overall trend across all MC sites (Figure 2C). A strong correlation between chromatin accessibility and editing frequency was observed, with highly accessible sites (MC sites 8, 7, and 4) displaying editing rates ranging from 8.17% to 29.21%, whereas low-accessibility sites (MC sites 1, 2, 3, 5, and 9) exhibited reduced editing rates ranging from 1.92% to 3.87% (Figure 2C). These findings confirm that AgroGem preserves endogenous chromatin context effects on CRISPR-Cas9 editing frequency.

Given that AgroGem maintains chromatin-dependent editing behavior, we next applied this platform to evaluate chromatin impacts on additional CRISPR-Cas systems. Notably, the MC sites targeted by CRISPR-Cas9 overlap with sequences that can also be recognized by *Lb*Cas12a, which requires a distinct 5’-TTTC-3’ protospacer-adjacent motif (PAM) located 5′ of the target site (Zhang et al., 2023); Figure 2C). Because *Lb*Cas12a is 11% smaller than *Sp*Cas9 in size, we evaluated whether its editing efficiency would be similarly influenced by chromatin accessibility (Zhang et al., 2023); Figure 2C). To address this, we constructed geminiviral replicon T-DNA vectors expressing *Lb*Cas12a and the corresponding CRISPR RNA targeting the same MC regions. Using AgroGem, Cas12a-mediated editing followed a similar chromatin-dependent pattern, with the highest editing frequency observed at MC site 8 (21.4%) and the lowest at MC site 5 (2.08%). Editing frequencies across all MC sites were highly correlated between *Sp*Cas9 and *Lb*Cas12a, with a Pearson correlation coefficient of 0.99 (*p* value = 6.6 x 10^-7^; Figure 2C). Together, these results establish AgroGem as a robust platform for the functional evaluation of genome editing reagents and demonstrate that chromatin context exerts a conserved influence on editing frequency across distinct CRISPR-Cas systems.

### AgroGem for rapid genetic dissection of DNA repair genes

To demonstrate the use of AgroGem for rapid functional validation of gene function, we characterized nine core genes involved in the non-homologous end joining (NHEJ) DNA repair pathway, including both canonical NHEJ (c-NHEJ) and alternative NHEJ (a-NHEJ) (Figure 3A), as well as one gene, *RECQ2*, involved in homologous recombination-associated DNA end resection and genome stability maintenance (Dorn and Puchta, 2019). The NHEJ-associated genes function at multiple stages of DNA repair, including DNA damage signaling (*ATM* and *ATR*), DSB recognition (*KU70* and *KU80*), end processing (*DNA Pol λ*, and *DNA Pol θ* also known as *DNA PolQ*), and ligation (*LIG4*, *XRCC4*, and *XRCC1*) (Figure 3A; Charbonnel et al., 2010; Wei et al., 2012; Chang et al., 2017; Hussmann et al., 2021a; Weiss et al., 2024; Frit et al., 2025). Homozygous T-DNA insertion mutants were obtained from the Arabidopsis Biological Resource Center (ABRC), prioritizing lines previously characterized in the literature (Supplemental Figure S2; Supplemental Table S1). Seeds from each mutant line were germinated and subjected to AgroGem-mediated transient transformation using a CRISPR-Cas9 construct targeting the *CHLI2* locus. CRISPR-induced mutation outcomes were then quantified by next-generation sequencing (Figure 3B).

**Figure 3.**
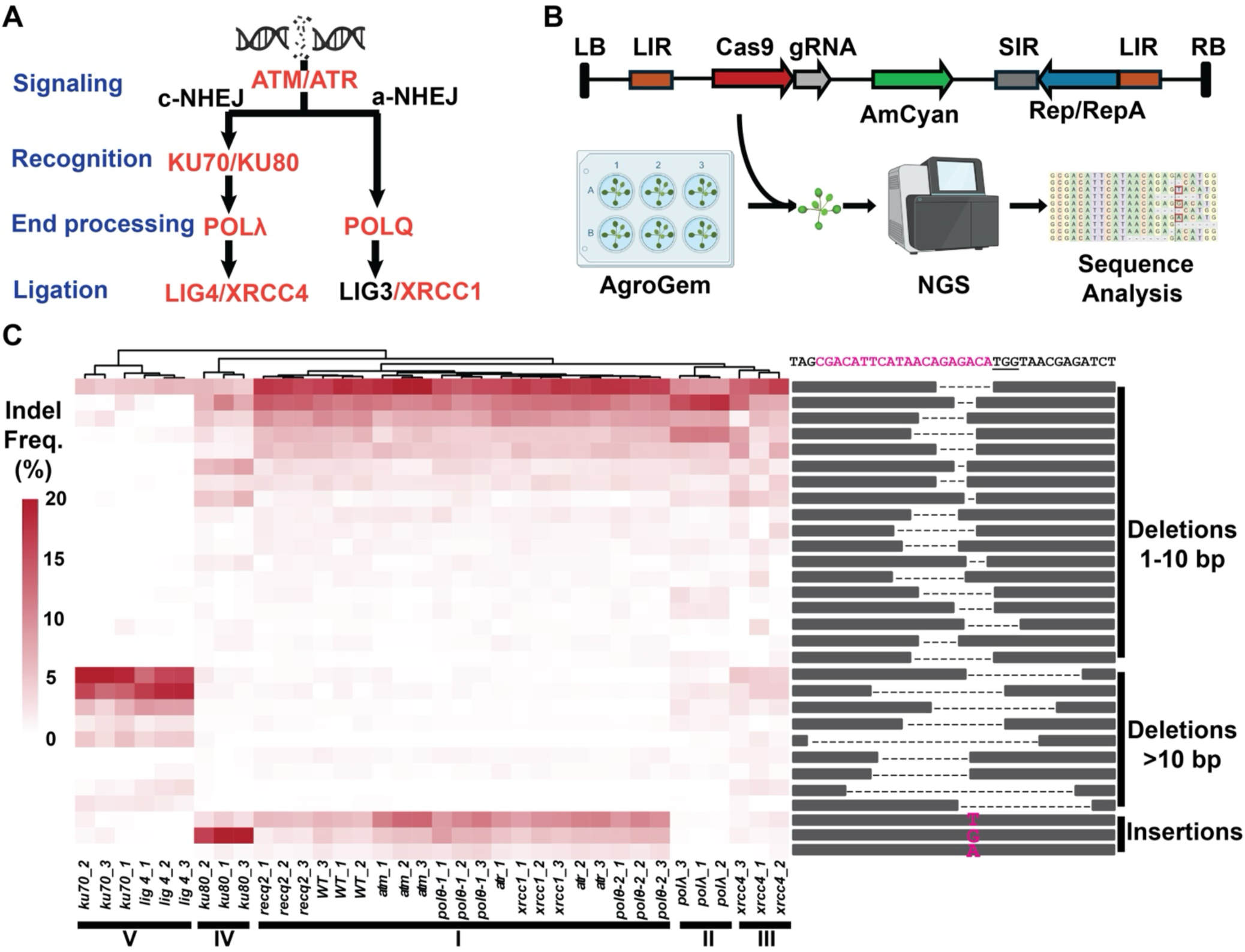
AgroGem enables high-resolution genetic dissection of NHEJ DNA repair pathways in Arabidopsis. (A) Schematic overview of key genes involved in canonical (c-NHEJ) and alternative (a-NHEJ) DNA repair pathways analyzed in this study, including DNA damage signaling, double-strand break recognition, end processing, and ligation. (B) Experimental workflow for mutation profiling using AgroGem. A geminiviral replicon T-DNA construct expressing Cas9, guide RNA, and the AmCyan reporter was delivered into Arabidopsis seedlings. Transformed tissues were collected, genomic DNA was extracted, and CRISPR-induced mutations were quantified by next-generation sequencing. LB and RB indicate T-DNA borders; LIR and SIR denote the long and short intergenic regions required for geminiviral replication; Rep/RepA represent viral replication proteins. (C) Hierarchical clustering of CRISPR-induced mutation spectra across DNA repair mutant lines. The heatmap shows normalized frequencies of top 30 mutation types across biological replicates. Mutants segregated into five major clusters (I–V) based on distinct repair signatures. Representative mutation categories are shown at right, including 1–10 bp deletions, >10 bp deletions, and 1-bp insertions. In deletion classes, dashed lines indicate deleted nucleotides, while in insertion classes, individual nucleotides denote the identity of the inserted base. The source data are provided in the Supplemental Source Data file S1.

To assess the impact of individual genes on NHEJ repair outcomes, we performed hierarchical clustering analysis based on indel mutation frequencies across all mutant lines (Figure 3C). For each sample, the frequency of each mutation type was normalized to its proportion of total edited reads. The top 30 most abundant mutation types were selected and grouped into three categories: small deletions (≤10 bp), large deletions (>10 bp), and single-base insertions (1 bp insertions). Notably, the three biological replicates for each mutant line clustered tightly together, demonstrating the high reproducibility of AgroGem. This analysis further uncovered distinct, genotype-specific repair signatures associated with individual NHEJ components. Based on mutation profile clustering (Figure 3C), the mutant lines segregated into five major groups. Cluster I displayed mutation patterns similar to wild type, whereas Clusters II–V exhibited distinct genotype-specific signatures. Cluster II was represented by *DNA Pol λ*; Cluster III by *xrcc4*; Cluster IV by *ku80*; and Cluster V by *lig4* and *ku70*. Cluster I included *atm*, *atr*, *xrcc1*, *DNA Pol θ*, and *recq2*, suggesting that disruption of these genes has a limited impact on the overall distribution of CRISPR-induced repair outcomes at this locus. The mutation profiles observed in Clusters II and V were consistent with previous reports showing that DNA Pol λ is primarily responsible for 1 bp insertion events during NHEJ, and loss of *lig4* or *ku70* increases the frequency of large deletions (Qi et al., 2013).

Prior studies suggested that the *ku80* mutant phenocopies *ku70* in CRISPR-induced mutation profiles, consistent with the obligate heterodimeric function of *KU70* and *KU80* (Qi et al., 2013; Shen et al., 2016). However, our high-resolution mutation profiling revealed that the repair signature of *ku80* (Cluster IV) differed markedly from that of *ku70*, particularly in the lower frequency of large deletions. Instead, the frequency of 1-bp insertions in the *ku80* mutant increased by 125% relative to wild type, whereas 1-bp insertions were nearly undetectable in the *ku70* mutant. Moreover, the 1-bp insertion spectrum in *ku80* differed from that of wild type, showing a pronounced enrichment for G insertions, whereas T insertions predominated in wild type (Figure 3C; Supplemental Figure S3). These findings suggest that KU80 may play a distinct and previously underappreciated role within the KU70/KU80 heterodimer in modulating DSB repair outcomes.

The *xrcc4* mutant in Cluster III also exhibited a distinct repair signature. Compared with wild type, *xrcc4* showed increased frequencies of both small and large deletions, accompanied by a marked reduction in 1-bp insertions. Because XRCC4 forms a stable complex with DNA Ligase IV (LIG4) and functions in the final ligation step of canonical NHEJ, a mutation profile similar to *lig4* would be anticipated (West et al., 2000). However, the mutation spectrum of *xrcc4* differed from that of the *lig4* mutant, particularly in the lower frequency of large deletions observed in *xrcc4*. These findings indicate that XRCC4 modulates the balance between deletion and insertion formation during NHEJ in a manner that is distinct from LIG4. Together, these results demonstrate that AgroGem enables rapid, high-resolution genetic dissection of DNA repair pathways and captures genotype-specific repair signatures in a quantitative and reproducible manner.

### AgroGem for assessing protein-protein interactions

Fluorescence reporter-based assays, such as bimolecular fluorescence complementation (BiFC), are widely used to interrogate protein-protein interactions (PPIs) for functional analysis in plants (Stefano et al., 2015). Efficient transient expression is essential for PPI assays. *Agrobacterium*-mediated delivery or protoplast transfection are the most used approaches (Xing et al., 2016). However, relatively low and variable transformation efficiencies have limited the routine application of *Agrobacterium*-mediated transient PPI assays in Arabidopsis. To determine whether AgroGem supports robust PPI assays in Arabidopsis, we selected two transcription factors, NUCLEAR FACTOR Y, Subunit B2 (NF-YB2) and NUCLEAR FACTOR Y, Subunit C3 (NF-YC3), previously shown to directly interact through their histone fold domains (Hackenberg et al., 2012). Three T-DNA constructs were generated: (1) wild-type coding sequences of the two proteins fused to complementary split GFP fragments, (2) an interaction-compromised variant carrying a mutation (E65R at NF-YB2) expected to weaken PPI (Siriwardana et al., 2016), and (3) a negative control construct lacking NF-YB2 sequences (Figure 4A). Each construct was introduced into Arabidopsis leaves using the AgroGem approach. Strong GFP complementation signals were detected in samples expressing the wild-type NF-YB2 and YC3 pair and co-localized with the DsRed marker, whereas GFP signals were barely detected in samples expressing the mutant protein construct or the empty vector control (Figure 4B). Together, these results demonstrate that AgroGem provides a robust and efficient transient expression platform for BiFC-based detection of PPIs in Arabidopsis.

**Figure 4.**
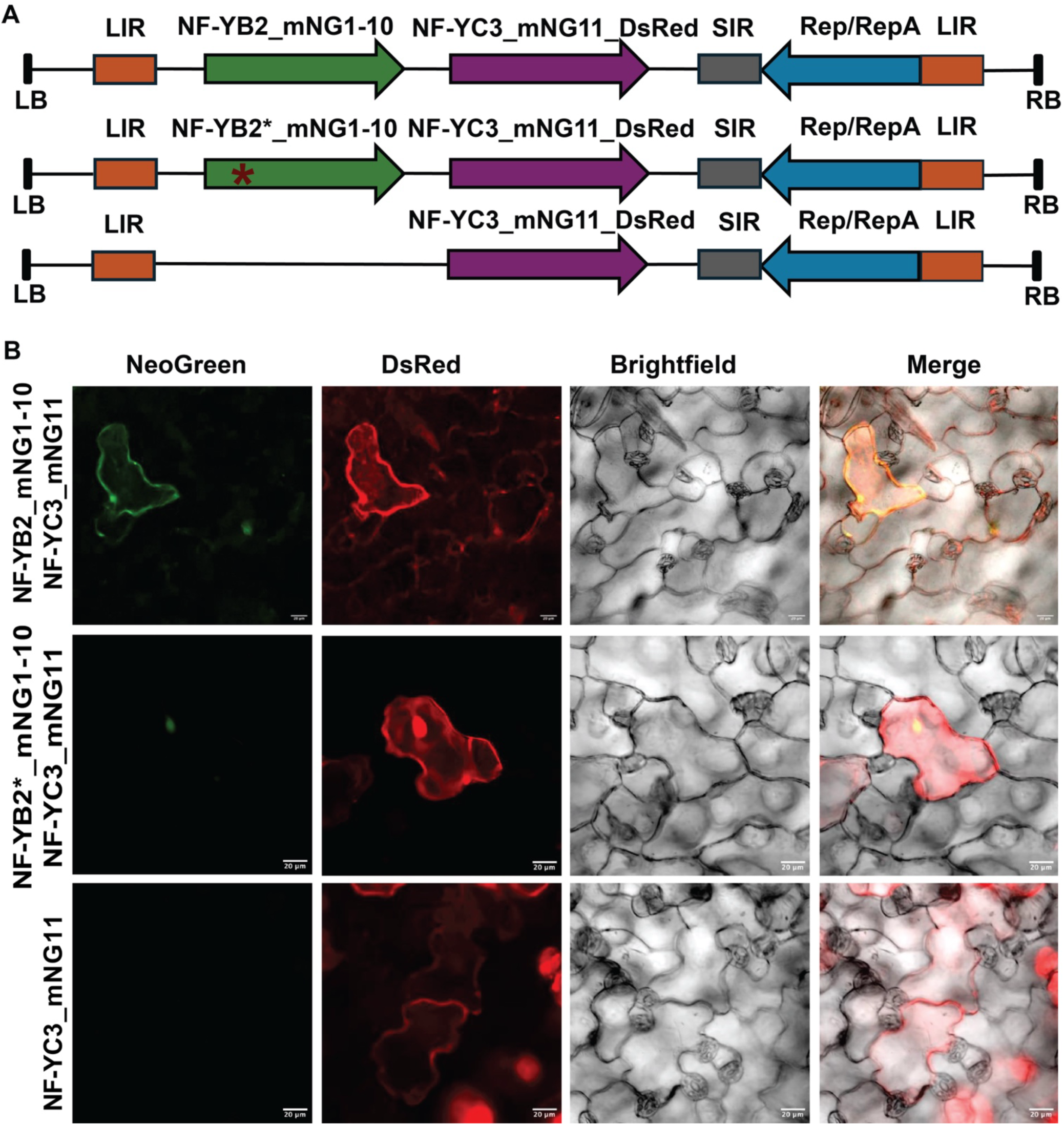
AgroGem supports bimolecular fluorescence complementation (BiFC) analysis in Arabidopsis. (A) Schematic representation of the geminiviral replicon T-DNA constructs used for BiFC assays. The wild-type construct expresses NF-YB2 fused to the N-terminal fragment of mNeonGreen (mNG1–10) and NF-YC3 fused to the complementary C-terminal fragment (mNG11), together with a DsRed reporter. An interaction-compromised mutant carrying the E65R substitution in NF-YB2 was generated as a negative control (NF-YB2*). An additional control construct expressing NF-YC3_mNG11_DsRed alone was included. LB and RB indicate T-DNA borders; LIR and SIR represent geminiviral replication elements; Rep/RepA denotes replicase proteins. (B) Representative fluorescence images of Arabidopsis epidermal cells following AgroGem-mediated transient expression. Strong NeoGreen fluorescence was observed in cells co-expressing wild-type NF-YB2_mNG1–10 and NF-YC3_mNG11, indicating protein–protein interaction. Minimal or no NeoGreen signal was detected in the E65R mutant (NF-YB2*) or NF-YC3-only control. DsRed marks transformed cells. Brightfield and merged images are shown. Scale bars: 20 μm.

### Scalable AgroGem enables efficient transient transformation across Brassicaceae species

A major limitation of existing plant transient transformation systems is limited throughput and poor scalability. To test whether AgroGem can be readily scaled, we germinated 1 - 2 Arabidopsis seeds per well in a 24-well plate and performed AgroGem transformation using the GVR T-DNA construct carrying the AmCyan reporter. Robust AmCyan fluorescence was detected in each well, indicating that the AgroGem workflow can be readily scaled to 24 samples processed in parallel. We next evaluated whether AgroGem is transferable to other *Brassicaceae* species. Seeds of camelina (*Camelina sativa*), pennycress (*Thlaspi arvense*), and *Brassica rapa* were germinated in 6 well plates, and the AgroGem protocol was applied using the same reporter construct. AmCyan fluorescence was consistently detected in each species, with signals observed in both cotyledons and true leaves (Figure 5). Thus, these results demonstrate that AgroGem is compatible with plate-based scaling and supports efficient transient transformation across multiple *Brassicaceae* species.

**Figure 5.**
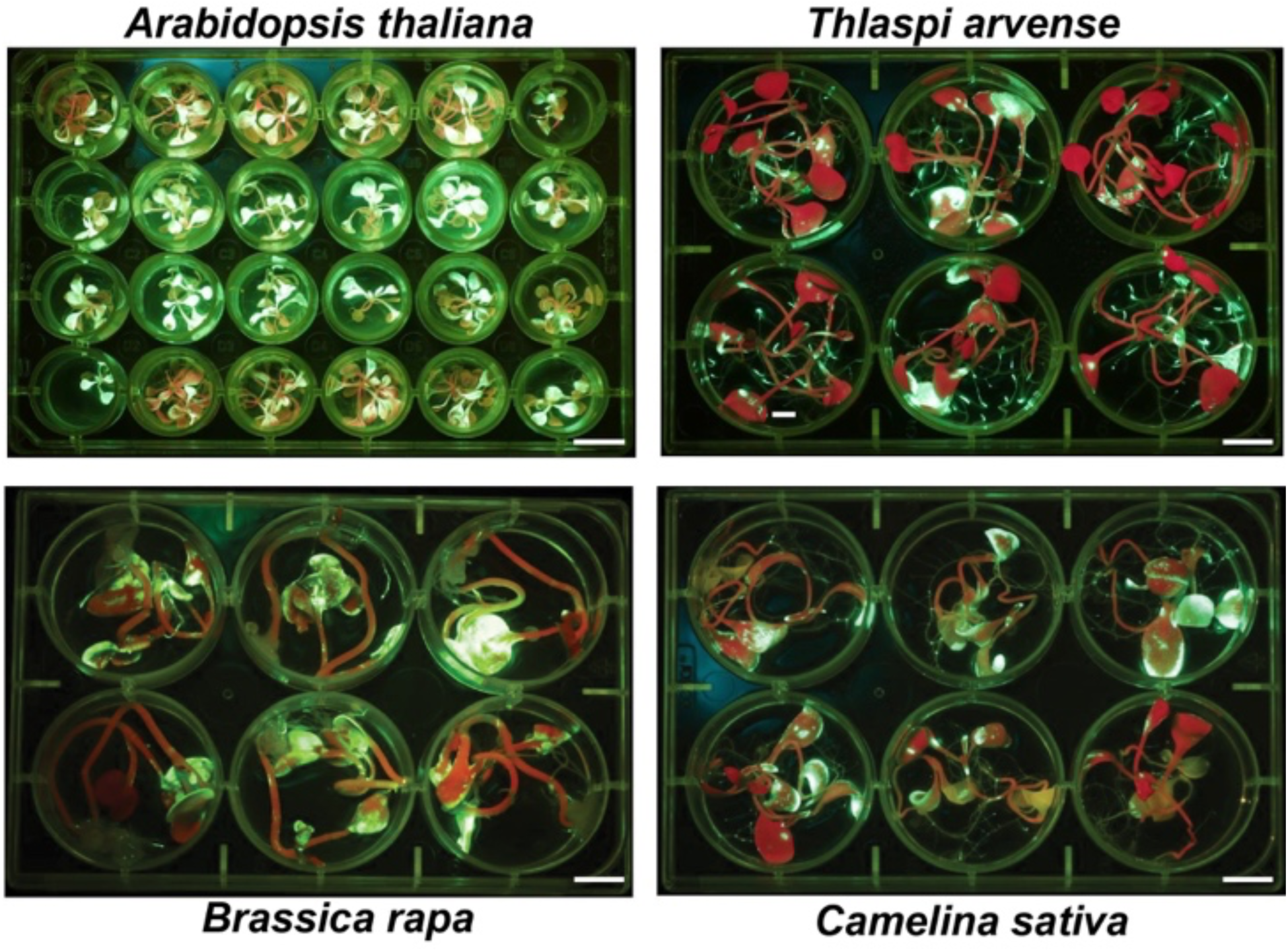
AgroGem enables scalable transient transformation across multiple *Brassicaceae* species. Representative fluorescence images showing AgroGem-mediated transient expression of the AmCyan reporter in seedlings of *Arabidopsis thaliana* (Col-0), *Thlaspi arvense* (var. mn106), *Brassica rapa* (var. R500), and *Camelina sativa* (var. Suneson). Seedlings were germinated in multiwell plates and subjected to the AgroGem protocol. Robust fluorescence signals were detected in cotyledons and true leaves across all species, demonstrating the scalability and cross-species applicability of the platform. Scale bars: 1 cm.

## Discussion

In this study, we developed AgroGem as an improved Agrobacterium-mediated transient transformation system that enables efficient and reproducible transgene delivery in Arabidopsis and related *Brassicaceae* species. AgroGem offers practical advantages over previously reported transient systems by reducing hands-on time and enabling straightforward scaling to plate-based formats. It is critical for applications that require parallel testing across many constructs or genotypes. Systematic optimization identified three key parameters that collectively produced the highest transformation efficiencies: use of a geminiviral replicon vector (GVR), optimization of commercially available Agrobacterium strains, and co-cultivation conditions. Geminiviral replicons are delivered as T-DNA and can be released and amplified in plant nuclei through rolling-circle replication, increasing transgene copy number and boosting expression (Gleba et al., 2004; Baltes et al., 2014). In addition, Agrobacterium-mediated transformation depends on both bacterial virulence and host permissiveness (Pitzschke, 2013). EHA105 is derived from the hypervirulent strain A281 and carries the pTiBo542 virulence region, which enhances T-DNA processing and transfer relative to commonly used strains such as GV3101 and C58C1 (Hood et al., 1993; Gelvin, 2003). This hypervirulence likely accounts for the superior performance of EHA105 observed in AgroGem. Dark co-cultivation can further promote infection by increasing bacterial motility, adhesion to plant tissues, and virulence gene activity (Oberpichler et al., 2008). From the host perspective, light-dependent immune outputs such as reactive oxygen species can restrict Agrobacterium infection (Pitzschke, 2013; Breen et al., 2023). Therefore, dark co-cultivation likely enhances transformation efficiency by both promoting bacterial infection and reducing host immune constraints. In this study, we focused only on commercially available Agrobacterium strains to ensure broad accessibility of the AgroGem method without the need for specialized strain exchange. As efforts to engineer next-generation hypervirulent Agrobacterium strains continue to advance, future evaluation of these strains may further improve the transformation efficiency and versatility of AgroGem (Goralogia et al., 2025).

One key motivation for developing AgroGem was to enable rapid and quantitative evaluation of genome editing reagents in intact tissues. Compared with AGROBEST and protoplast-based assays, AgroGem produced substantially higher editing rates at the tested locus in this study, providing a practical advantage for benchmarking editing systems with quantitative readouts. In addition, the current AGROBEST method relies on the immune-deficient *efr1* (EF-Tu Receptor 1) mutant background to enhance transformation efficiency (Wu et al., 2014). *EFR1* encodes a leucine-rich repeat receptor-like kinase involved in the recognition of bacterial EF-Tu and activation of pattern-triggered immunity during Agrobacterium infection (Zipfel et al., 2006). Consequently, use of the *efr1* mutant background could introduce confounding effects associated with altered immune signaling. In contrast, AgroGem achieved robust transient expression directly in the wild-type Col-0 background, thus avoiding perturbation of endogenous immune pathways.

In addition to improving transient transformation and transgene expression efficiency, AgroGem generated mutation profiles that closely resembled those obtained through stable floral dip transformation. This capability is particularly important because genome editing outcomes are strongly influenced by chromatin accessibility and local genomic context. Using multi-copy target sites spanning distinct chromatin environments, we demonstrated that AgroGem preserves chromatin-dependent effects on editing efficiency and that these effects are conserved across different CRISPR-Cas systems. Together, these findings establish AgroGem as a rapid and efficient alternative to protoplast-based assays and stable transformation for evaluating genome editing reagents. This workflow should be readily adaptable to additional genome editors, including base editors, prime editors, and emerging RNA-guided transposases and recombinases (Capdeville et al., 2023; Vats et al., 2024; Makarova et al., 2025; Perry et al., 2025).

AgroGem also enables us to rapidly dissect DNA repair pathways through quantitative profiling of CRISPR-induced repair outcomes across genetic backgrounds. Although many DNA repair genes have been individually characterized in plants, functional assignments have largely relied on homology to yeast and mammalian systems (Puchta, 2005). In budding yeast (*Saccharomyces cerevisiae*), classical genetic studies established the core architecture of non-homologous end joining (NHEJ), identifying key roles for Ku70/Ku80, DNA ligase IV, and associated factors in end processing and ligation (Symington and Gautier, 2011). More recently, high-resolution CRISPR-based profiling in human cells has systematically quantified mutation spectra across repair gene knockouts, revealing pathway-specific signatures and functional hierarchies (Hussmann et al., 2021b; de Alba et al., 2025). Comparable systematic and quantitative analyses, however, have been largely lacking in plants. Using a panel of core NHEJ mutants, we found that hierarchical clustering of mutation spectra segregated mutants into distinct groups, demonstrating that AgroGem can resolve pathway-specific repair signatures in planta. Several patterns were consistent with conserved roles observed in yeast and mammalian systems. For example, DNA Pol λ clustered separately and was associated with elevated 1-bp insertions, consistent with its conserved function in gap filling and templated insertion during NHEJ (Weiss et al., 2024). Likewise, loss of *Ku70* or *Lig4* increased the frequency of large deletions, reflecting their central roles in DNA end protection and ligation, respectively (Zhang et al., 2010). These results support substantial evolutionary conservation of core NHEJ mechanisms across eukaryotes.

Having established this conserved framework, AgroGem also uncovered repair signatures that diverge from expectations based on yeast and mammalian systems. In budding yeast and mammalian cells, KU70 and KU80 function as an obligate heterodimer, and loss of either subunit typically produces highly similar NHEJ phenotypes (Hussmann et al., 2021b). However, in our study, *Ku80* displayed a mutation spectrum distinct from *Ku70*, characterized by reduced large deletions and increased 1-bp insertions with a distinct nucleotide preference. This divergence suggests that, although the KU complex is structurally conserved, KU70 and KU80 may contribute non-equivalently to repair outcome regulation in plants. Similarly, while XRCC4 and LIG4 act together in canonical NHEJ in both yeast and mammals, the mutation profile of *Xrcc4* differed quantitatively from that of *Lig4*, indicating that XRCC4 may influence DSB repair outcomes beyond simply stabilizing LIG4-mediated ligation. It is important to note that T-DNA insertion alleles may not represent complete loss-of-function mutations and can produce partial loss-of-function or unexpected phenotypes (Jupe et al., 2019; Song et al., 2026). Nevertheless, the reproducibility and resolution of the AgroGem approach enabled detection of repair signatures that would be difficult to identify using conventional approaches. Together, these findings demonstrate that AgroGem provides a powerful framework for high-resolution analysis of plant DNA repair pathways, revealing both conserved repair mechanisms and potentially lineage-specific features of NHEJ regulation that warrant further genetic and biochemical investigation.

Beyond genome editing and repair profiling, AgroGem provides a practical transient expression platform for other functional assays. BiFC in particular is widely performed in *N. benthamiana* because of robust leaf agroinfiltration, yet this approach is difficult to scale and lacks the extensive genetic resources available in Arabidopsis (Walter et al., 2004). Using NF-YB2 and NF-YC3 as a test pair, we showed that AgroGem supports robust BiFC readouts in Arabidopsis leaves, demonstrating broader utility for PPI testing in the native Arabidopsis context. Finally, AgroGem was readily extended to multiple *Brassicaceae* species, including camelina, pennycress, and *Brassica rapa*, supporting its portability and potential as a scalable transient platform beyond Arabidopsis.

In summary, AgroGem provides a rapid, efficient, and scalable Agrobacterium-mediated transient transformation system that supports quantitative functional genetics in plants. By enabling high-efficiency transgene delivery in intact tissues while preserving endogenous chromatin and DNA repair contexts, AgroGem facilitates scalable evaluation of genome editing strategies and DNA repair mechanisms. The platform can be readily adapted for broader applications, including systematic pathway mapping and protein characterization. Although extension to additional plant species may require optimization of Agrobacterium strains, co-cultivation conditions, or vector architecture, AgroGem establishes a flexible framework that combines the speed and scalability of transient expression systems with the biological relevance of native tissue contexts. Furthermore, its compatibility with plate-based formats enables integration with automation platforms, laying the foundation for high-throughput functional genomics workflows.

## Materials and methods

### T-DNA vector construction

The CRISPR/Cas9 construct was created using the Golden Gate assembly method as outlined previously (Čermák et al., 2017). The gRNA sequences targeting the multicopy CRISPR site (MC site) and the CHLI2 locus were first cloned into the pMOD_B2301 entry vector. Final T-DNA constructs were assembled from the AtUbi10-driven Cas9 expression cassette (pMOD_A0102), the corresponding gRNA modules targeting MC site and CHLI2 (Weiss et al., 2022), an AmCyan fluorescent reporter driven by the *Sl*Ubi promoter (Addgene #197731; Chamness et al., 2024), and the geminiviral replicon T-DNA backbone pTRANS_221 (Addgene #91115; Čermák et al., 2017). The resulting construct enabled simultaneous expression of Cas9, gRNA, and the AmCyan reporter following Agrobacterium-mediated delivery.

### Seed sterilization and growth conditions

Seeds were surface sterilized by incubation in 1 mL of 70% ethanol for 5 minutes with vortexing. The seeds were briefly centrifuged to remove ethanol and then incubated in 1 mL of 30% (v/v) commercial bleach supplemented with 1 μL Triton X-100 for 5 minutes with vortexing. Following sterilization, the seeds were collected by brief centrifugation, and then washed three times with 1 mL sterile water. Each wash was performed by vortexing for 5 minutes, followed by brief centrifugation and removal of the supernatant. After sterilization, seeds were suspended in 1 mL sterile one-half-strength Murashige and Skoog (½ MS) medium supplemented with Gamborg vitamins and stratified at 4 °C for 3–4 d. Following stratification, six seeds were transferred to each well of a 6-well plate containing 2 mL of ½ MS medium supplemented with 0.5% sucrose (pH 5.5). Plates were sealed with surgical tape and incubated in a growth chamber at 24 °C under a 16 h light/8 h dark photoperiod until use.

### AGROBEST and AgroGem transformation

The AGROBEST protocol was performed as previously described using either one- or two-week-old Arabidopsis seedlings (Wu et al., 2014). For AgroGem transformation, Arabidopsis seedlings were grown in 6-well plates for 14 days prior to inoculation. To prepare Agrobacterium cultures, a single colony of *Agrobacterium tumefaciens* strain (all three agrobacterium strains, EHA105, GV3101 and C58C1, are purchased from Gold Biotechnology Inc., St. Louis, MO, U.S.A.). carrying T-DNA plasmid was inoculated into 5 mL LB medium supplemented with the appropriate antibiotics and incubated overnight at 30°C with shaking. Cells were collected by centrifugation and resuspended in 5 mL AB-MES medium (17.2 mM K₂HPO₄, 8.3 mM NaH₂PO₄, 18.7 mM NH₄Cl, 2 mM KCl, 1.25 mM MgSO₄, 100 μM CaCl₂, 10 μM FeSO₄, 50 mM MES, and 2% [w/v] glucose, pH 5.5) supplemented with 200 μM acetosyringone. Cultures were incubated overnight at 30°C with shaking. OD_600_ of the culture was measured and diluted with ½ AB-MES ¼ MS (mixing AB-MES with ½ MS as 1:1, pH 5.5) with 200 uM acetosyringone to adjust OD_600_ to 0.02. Growth medium was then removed from each well containing seedlings and replaced with 2 mL of the Agrobacterium culture. Plates were wrapped in aluminum foil and incubated at 22 °C for 3 days under continuous darkness for co-cultivation. Following co-cultivation, the inoculation medium was removed and replaced with 1 mL of ½ MS medium supplemented with 0.5% sucrose (pH 5.5) and 250 μg/mL timentin. Plates were wrapped in aluminum foil and incubated for an additional 2-3 days at 24 °C to allow recovery and transgene expression. Leaves exhibiting AmCyan fluorescence were subsequently collected for downstream analyses.

### Protoplast isolation and transfection

Protoplast isolation and transfection were performed using a modified version of the PEG-mediated method (Yoo et al., 2007; Li et al., 2016). In brief, leaves from 14-day young Arabidopsis seedlings were sliced into small pieces with a razor blade and digested with the enzyme solution (1.5% Cellulase, 0.75% Macerozyme, Kanematsu USA Inc.) for 4-5 hours on a shaker at 40 rpm. The digested tissues were filtered through a 70 M nylon filter (Fisher Scientific, Waltham, Massachusetts, USA) into W5 buffer (2 mM MES with pH5.7, 154 mM NaCl, 125 mM CaCl2, 5 mM KCl). Protoplasts were collected and resuspended in W5 buffer with a gentle centrifuge at 100xg for 5 minutes. The number of protoplasts was estimated using a hemocytometer. Roughly 200,000 protoplasts were mixed with DNA plasmids (15 g per construct) in 20% PEG buffer and incubated at room temperature in the dark for 48 hours. Transformation efficiencies were monitored by transforming protoplasts with a plasmid encoding GFP. Three transformation replicates were performed for each experiment. After 48-hour incubation, transformed protoplasts from all replicates were pooled to extract genomic DNA (Li et al., 2016).

### Genomic DNA Extraction

Leaf tissues were collected for genomic DNA extraction using a modified SDS-based method (Edwards et al., 1991). Briefly, samples were homogenized with metal beads in the extraction buffer (200 mM Tris-HCl, 250 mM NaCl, 25 mM EDTA, and 0.5% SDS, pH 8.0). Genomic DNA was recovered by isopropanol precipitation, washed with 75% ethanol, air-dried, and resuspended in 50 μL of 10 mM Tris-HCl (pH 8.0) or nuclease-free water.

### Characterization of T-DNA mutant lines

Homozygous T-DNA insertion lines used in this study were obtained from the Arabidopsis Biological Resource Center (ABRC). T-DNA insertion sites and gene-specific primers were identified using the SIGnAL T-DNA Express database (http://signal.salk.edu/cgi-bin/tdnaexpress; (O’Malley et al., 2015)). For each line, two gene-specific primers flanking the predicted T-DNA insertion site were designed (Supplemental Table S2). Genotypes were confirmed by PCR using the gene-specific primers together with the T-DNA left border primer LBb1.3 (ATTTTGCCGATTTCGGAAC) as previously described (O’Malley et al., 2015). Homozygous insertion lines were identified based on the absence of wild-type alleles and the presence of T-DNA-specific amplification products.

### Mutation genotyping and characterization of mutation profiles

Genome editing events were initially assessed by PCR amplification of the *CHLI2* and MC sites target loci, followed by cleaved amplified polymorphic sequence (CAPS) analysis. PCR was performed using GoTaq Green Master Mix (Promega, Madison, WI, USA) according to the manufacturer’s instructions with an annealing temperature of 55°C. Primer sequences are provided in Supplemental Table S2. PCR amplicons were digested with BsmAI (*CHLI2*) or AluI (MC site). For quantitative analysis of editing outcomes, target amplicons were subjected to Illumina paired-end sequencing (Genewiz, South Plainfield, NJ, USA). Sequencing reads were analyzed using CRISPResso2 to quantify indel frequencies and characterize mutation profiles (Clement et al., 2019). For mutation spectrum analysis, indel-containing reads were extracted from CRISPResso2 output files, and mutation types exceeding a 2% frequency threshold were retained for downstream analysis. Data processing, visualization, and hierarchical clustering were performed in R Studio (v4.1.0) using the ggplot2 package (Wickham, 2009).

### Luciferase assay

Leaves exhibiting AmCyan fluorescence were collected following the AgroGem protocol. Approximately 10 leaves were harvested per sample, and fresh tissue weight was recorded for normalization. Luciferase activity was quantified using the Bio-Glo Luciferase Assay System (Promega, Madison, WI, USA) according to the manufacturer’s instructions.

### Bimolecular fluorescence complementation (BiFC) assay

BiFC assays were performed by transient expression in Arabidopsis through the AgroGem protocol. Coding sequences were cloned in-frame with N- or C-terminal fragments of the split mNeonGreen2 (mNG2) fluorescent protein in AgroGem T-DNA vectors. Plants were allowed to recover for 3 days after AgroGem treatment prior to confocal microscopy. Excised leaves were imaged on a Nikon AX R Confocal microscope with a 20X water immersion objective (NA 0.95, RI 1.333), using separate laser lines and detectors for mNG2 (488 ex, 507-551 em) and DsRed (561 ex, 593-625 em). Transmitted light was captured through the 561 laser line. All images were collected in parallel and without alteration of any excitation or detection parameters.

### Statistical analysis

Pearson correlation coefficients were calculated in R v4.1.0 to assess the similarity between indel frequency distributions generated by AgroGem and floral dip transformation in the Col-0 and *lig4* backgrounds (Figure 2B), and to compare editing frequencies between *Sp*Cas9 and *Lb*Cas12a across all MC sites (Figure 2C). Unsupervised hierarchical clustering was performed using the pheatmap package in R Studio (v4.1.0). Samples were clustered based on Euclidean distance using complete linkage, as implemented in the pheatmap function (Figure 3C). Statistical analyses in Figure 1B and 2A were performed using a one-tailed, two-sample Student’s t-test assuming equal variances.

### Accession numbers

NGS raw data can be found at NCBI Sequence Read Archive under PRJNA1442664. All CRISPResso and data analysis code can be found here: https://github.com/ZhangLab-UMN/AgroGem_manuscript

## Acknowledgements

J.K. and F.Z. is supported by the National Science Foundation (IOS-2040218 and IOS-2206920) awards. S.G. is supported by the Non-Assistance Cooperative Agreement with USDA-ARS (58-5062-3-017). K.G. is supported by the National Science Foundation (IOS-2029549 and DBI-2042159). We thank Dr. Savio De Siqueira Ferreira from Dr. Michael Smanski’s laboratory at the University of Minnesota for generously providing the Camelina seeds.

## Author Contributions

S.K., K.G. and F.Z. conceived and planned the study. S.G., O.S., Z.M., and J.K. performed the experiments. S.G., Z.M., K.G. and F.Z. analyzed the data. S.G., Z.M., S.K., K.G. and F.Z. wrote and edited the manuscript with input from all authors. Microscopy work was supported by the resources and staff at the University of Minnesota University Imaging Centers (UIC, SCR_020997).

## Supporting information

**Supplemental Figure S1.** Schematic map and sequences of the geminiviral replicon T-DNA vector.

**Supplemental Figure S2.** T-DNA insertion lines used for DNA repair pathway analysis.

**Supplemental Figure S3.** CRISPR-Cas9 mutation profiles in wild-type and *ku80* Arabidopsis plants.

**Supplemental Table S1.** Summary of T-DNA insertion lines for DNA repair pathway analysis.

**Supplemental Table S2.** Summary of synthesized oligo sequences.

**Supplemental File S1.** Source data file.

