## Supplemental information for "AgroGem: A Rapid and Scalable Transient Transformation System for Functional Genetics in Multiple Plant Species"

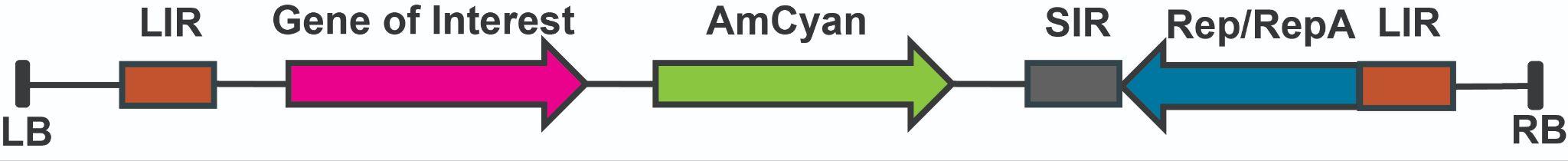

**LB**

tggcaggatatattgtggtgtaaacatgctccaccatgttggcaagctgctctagccaatacgcaaaccgcctctccccgcgcgttggccgattcattaatg

cagctggcacgacaggtttcccgactggaaagcgggcagtgagcgcaacgcaattaatgtgagttagctcactcattaggcaccccaggctttacacttt

atgcttccggctcgtatgttgtgtggaattgtgagcggataacaatttcacacaggaaacagctatgaccatgattacgaattcgagctcaaagtttaacgcg

**LIR**

ttagcagaaggcatgttgttgtgactccgaggggttgcctcaaactctatcttataaccggcgtggaggcatggaggcaggggtattttggtcattttaatag

atagtggaaaatgacgtggaatttacttaaagacgaagtctttgcgacaagggggggcccacgccgaatttaatattaccggcgtggcccccccttatcgc

gagtgctttagcacgagcggtccagatttaaagtagaaaatttcccgcccactagggttaaaggtgttcacactataaaagcatatacgatgtgatggtattt

gatggagcgtatattgtatcaggtatttccgttggatacgaattattcgtacgaccctcggtaccgatcggcgcgcccctgcaggtagatcgctcgtcgac

**Gene of interest**

NNNNNNNNNNNNNNNNNNNNNNNNNNNNNNNNNNNNNNNNNNNNNNNNNNNNNNNNNN

**SlUbi Promoter**

gagctctaccggggcgcgccagacataaattctttttttgtaatgctcaaataaattgagtaaaaaagaatgaaattgagtgatttttttttaatcataagaaaataaataattaatttcaatataataaaacagtaatataatttcataaatggaattcaatacttacctcttagatataaaaaataaataataaaaataaagtgtttctaataaacccgcaatttaaataaaatattataatattttcaatcaaatttaaataattatattaaaatatcgtagaaaaagagcaatatataatacaagaaagaagatttaagtacaattatcaactattattatactctaattttgttatattaatttcttacggttaaggtcatgttcacgataaaactcaaaatacgctgtatgaggacatattttaaattttaaccaataataaactaagttatttttagtatatttttttgtttaacgtgacttaatttttcttttctagaggagcgtgtaagtgtcaacctcattctcctaattttcccaaccacataaaaaaaaaataaaggtagcttttgcgtgttgatttggtacactacacgtcattattacacgtgttttcgtatgattggttaatccatgaggcggtttcctctagagtcggccataccatctataaaataaagctttctgcagctcattttttcatcttctatctgatttctattataatttctctgaattgccttcaaatttctctttcaaggttagaatttttctctattttttggtttttgtttgtttagattctgagtttagttaatcaggtgctgttaaagccctaaattttgagtttttttcggttgttttgatggaaaatacctaacaattgagttttttcatgttgttttgtcggagaatgcctacaattggagttcctttcgttgttttgatgagaaagcccctaatttgagtgtttttccgtcgatttgattttaaaggtttatattcgagtttttttcgtcggtttaatgagaaggcctaaaataggagtttttctggttgatttgactaaaaaagccatggaattttgtgtttttgatgtcgctttggttctcaaggcctaagatctgagtttctccggttgttttgatgaaaaagccctaaaattggagtttttatcttgtgttttaggttgttttaat

**AmCyan**

ccttataatttgagttttttcgttgttctgattgttgtttttatgaattttgcagaatggccctgtccaacaagttcatcggcgacgacatgaagatgacctaccacatggacggctgcgtgaacggccactacttcaccgtgaagggcgagggcagcggcaagccctacgagggcacccagacctccaccttcaaggtgaccatggccaacggcggccccctggccttctccttcgacatcctgtccaccgtgttcatgtacggcaaccgctgcttcaccgcctaccccaccagcatgcccgactacttcaagcaggccttccccgacggcatgtcctacgagagaaccttcacctacgaggacggcggcgtggccaccgccagctgggagatcagcctgaagggcaactgcttcgagcacaagtccaccttccacggcgtgaacttccccgccgacggccccgtgatggccaagatgaccaccggctgggacccctccttcgagaagatgaccgtgtgcgacggcatcttgaagggcgacgtgaccgccttcctgatgctgcagggcggcggcaactacagatgccagttccacacctcctacaagaccaagaagcccgtgaccatgccccccaaccacgcggtggagcaccgcatcgccagaaccgacctggacaagggcggcaacag

**SlUbi Terminator**

cgtgcagctcaccgagcacgccgtggcccacatcacctccgtggtgcccttctagcttgttgtggttgtctggttgcgtctgttgcccgttgtctgttgcccattgtggtggttgtgtttgtatgatggtcgttaaggatcatcaatgtgttttcgctttttgttccattctgtttctcatttgtgaataataatggtatctttatgaatatgcagtttgtggtttcttttctgattgcagttctgagcattttgtttttgcttccgtttactataccacttacagtttgcactaatttagttgatatgcgagccatctgatgtttgat

**SIR**

gattcaaatggcgtttatgtaactcgtacccgctgagctctcccggcgcgccgatatcgagctcagtgtttgatcgccggcggtaccgagtgtacttcaagtcagtgggaaatcaataaaatgattattttatgaatatatttcattgtgcaagtagatagaaattacatatgttacataacacacgaaataaacaaaaaaagacaatccaaaaacaaacaccccaaaaaaaataatcactttagataaactcgtatgaggagaggcacgttcagtgactcgacgattcccgagcaaaaaaagtctccccgtcacacatgtagtgggtgacgcaattatctttaaagtaatccttctgttgacttgtcattgataacatccagtcttcgtcaggattgcaaagaattataga

**Rep/RepA**

agggatcccaccttttattttcttcttttttccatatttagggttgacagtgaaatcagactggcaacctattaattgcttccacaatgggacgaacttgaaggggatgtcgtcgatgatattataggtggcgtgttcatcgtagttggtgaaatcgatggtaccgttccaatagttgtgtcgtccgagacttctagcccaggtggtctttccggtacgagttggtccgcagatgtagaggctggggtgtcggattccattccttccattgtccttgttaaatcggccatccattcaaggtcagattgagcttgttggtatgagacaggatgtatgtaagtataagcgtctatgcttacatggtatagatgggtttccctccaggagtgtagatcttcgtggcagcgaagatctgattctgtgaagggcgacacatacggttcaggttgtggagggaataatttgttggctgaatattccagccattgaagctttgttgcccattcatgagggaattcttccttgatcatgtcaagatattcctccttagacgttgcagtctggataatagttctccatcgtgcgtcagatttgcgaggagaaaccttatgatctcggaaatctcctctggttttaatatctccgtcctttgatatgtaatcaaggacttgtttagagtttctagctggctggatattagggtgatttccttcaaaatcgaaaaaagaaggatccctaatacaaggttttttatcaagctggagaagagcatgatagtgggtagtgccatcttgatgaagctcagaagcaacaccaaggaagaaaataagaaaaggtgtgagtttctcccagagaaactggaataaatcatctctttgagatgagcacttgggataggtaaggaaaacatatttagattggagtctgaagttcttacta

**LIR**

gcagaaggcatgttgttgtgactccgaggggttgcctcaaactctatcttataaccggcgtggaggcatggaggcaggggtattttggtcattttaatagatagtggaaaatgacgtggaatttacttaaagacgaagtctttgcgacaagggggggcccacgccgaatttaatattaccggcgtggcccccccttatcgcgagtgctttagcacgagcggtccagatttaaagtagaaaatttcccgcccactagggttaaaggtgttcacactataaaagcatatacgatgtgatggtatttgat

**RB**

ggagcgtatattgtatcaggtatttccgttggatacgaattattcgtacgaccctcatagtttaaactatcagtgtttgacaggatatattggcgggtaaacctaagagaaaagagcgttta

**Figure S1. Schematic map and sequences of the geminiviral replicon T-DNA vector.** Schematic representation of the geminiviral replicon T-DNA construct used for transient expression of genes of interest and the AmCyan reporter. DNA sequences corresponding to each vector component are shown. LB and RB denote the left and right T-DNA borders, respectively. LIR and SIR indicate the long and short intergenic regions required for geminiviral replication, and Rep/RepA encode the viral replication proteins that mediate replicon amplification in plant cells.

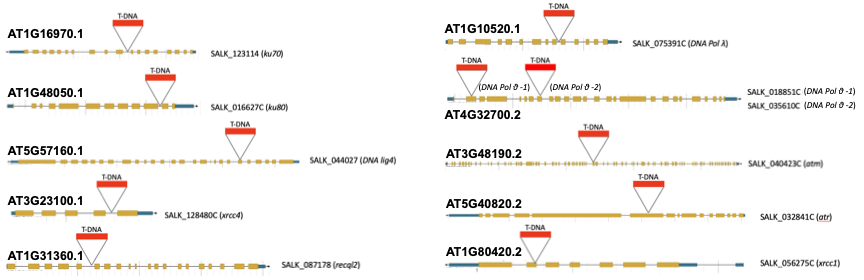

**Figure S2. T-DNA insertion lines used for DNA repair pathway analysis.** Schematic representation of the Arabidopsis T-DNA insertion lines analyzed in this study. Gene models are shown with exons represented by boxes and introns by connecting lines. Triangles indicate the positions of T-DNA insertions within each locus. Corresponding SALK accession numbers are shown for *ku70* (*At1g16970.1*), *ku80* (*At1g48050.1*), *lig4* (*At5g57160.1*), *xrcc4* (*At3g23100.1*), *recq2* (*At1g31360.1*), *polλ* (*At1g10520.1*), *polθ* (*At4g32700.2*), *atm* (*At3g48190.2*), *atr* (*At5g40820.2*), and *xrcc1* (*At1g80420.2*). Gene models and T-DNA insertion positions were mapped to the Arabidopsis TAIR10 reference genome.

**Figure S3**

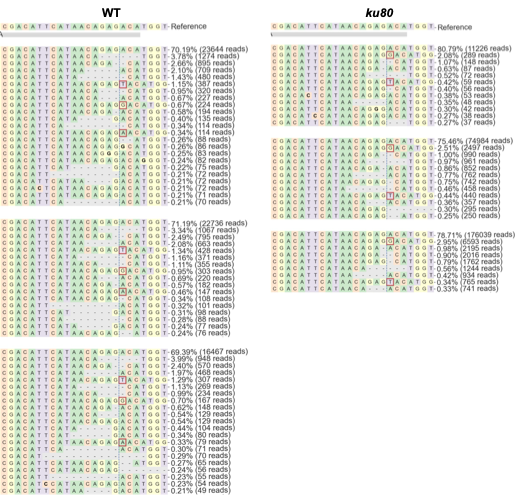

**Figure S3. CRISPR-Cas9 mutation profiles in wild-type and *ku80* Arabidopsis plants.** Representative CRISPR-Cas9-induced mutation profiles at the *CHLI2* target site in wild-type (Col-0) and *ku80* mutant backgrounds. Three biological replicates are shown for each genotype. The reference sequence is shown at the top of each panel with the 20-bp targeted site highlighted. Dashes indicate deleted nucleotides, and nucleotides highlighted by red boxes represent insertions. The vertical dashed line marks the predicted Cas9 cleavage site. For each mutation type, the percentage of total sequencing reads and the corresponding read count are indicated on the right.
